# Re-analysis of mobile mRNA datasets highlights challenges in the detection of mobile transcripts from short-read RNA-Seq data

**DOI:** 10.1101/2024.07.25.604588

**Authors:** Pirita Paajanen, Melissa Tomkins, Franziska Hoerbst, Ruth Veevers, Michelle Heeney, Hannah Rae Thomas, Federico Apelt, Eleftheria Saplaoura, Saurabh Gupta, Margaret Frank, Dirk Walther, Christine Faulkner, Julia Kehr, Friedrich Kragler, Richard J. Morris

## Abstract

Short-read RNA-Seq analyses of grafted plants have led to the proposal that large numbers of mRNAs move over long distances between plant tissues, acting as potential signals. The detection of transported transcripts by RNA-Seq is both experimentally and computationally challenging, requiring successful grafting, delicate harvesting, rigorous contamination controls and data processing approaches that can identify rare events in inherently noisy data. Here, we perform a meta-analysis of existing datasets and examine the associated bioinformatic pipelines. Our analysis reveals that technological noise, biological variation and incomplete genome assemblies give rise to features in the data that can distort the interpretation. Taking these considerations into account, we find that a substantial number of transcripts that are currently annotated as mobile are left without support from the available RNA-Seq data. Whilst several annotated mobile mRNAs have been validated, we cannot exclude that others may be false positives. The identified issues may also impact other RNA-Seq studies, in particular those using single nucleotide polymorphisms (SNPs) to detect variants.

## Introduction

Transport of messenger RNA (mobile mRNA) between cells and tissues (1–7) has been shown to play a role in several physiological and developmental processes in plants, such as tuberisation (8), leaf development (9) and meristem maintenance (10). For some cases, their influence on plant development has been described (9, 11–13), but for most mobile mRNAs the relevance of transport remains to be determined (14–17).

Grafting is an ancient technique that has been and is still used in traditional horticulture to confer abiotic and biotic resistance to crops (18). Grafting (18) has also become the method of choice for identifying mobile mRNAs as it reduces the risk of contamination compared to other approaches (16, 19), Figure 1. The pioneering work in the detection of mobile mRNA required technical expertise in these traditionally horticultural skills, as well as advanced knowledge of molecular biology and physiology. In early breakthroughs, important mobile signals in plants were discovered, such as florigen (20) of which Flowering Locus T protein (possibly also its mRNA (21, 22)) is now known to be a major contributing factor (23, 24) The use of grafting combined with advanced sequencing technologies and computational analyses has expanded the possibilities of the field to new heights.

**Fig. 1.**
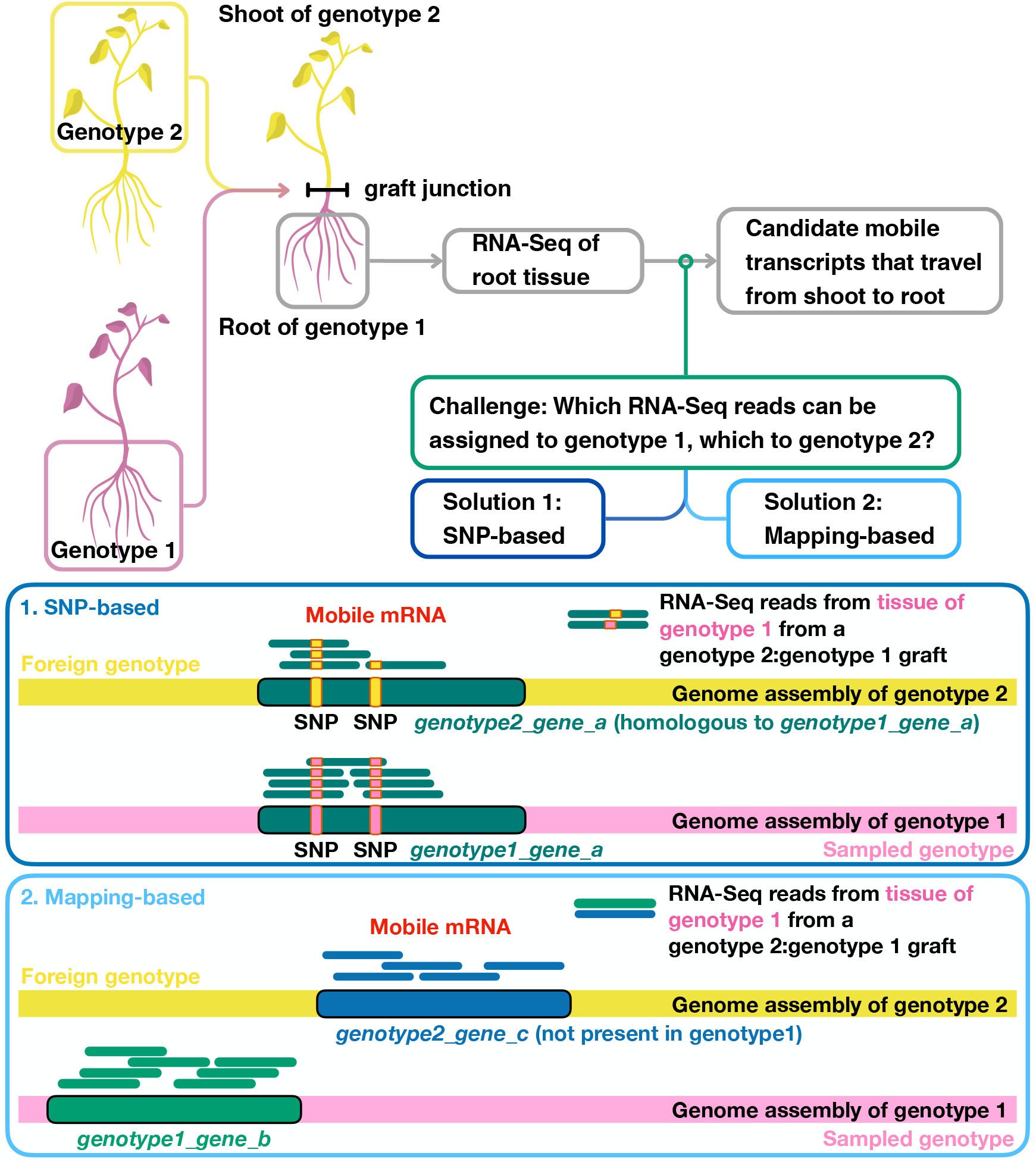
Grafting coupled with RNA-Seq has been employed to identify transcripts that move from tissue of one genotype/species/ecotype/cultivar into tissue of another genotype/species/ecotype/cultivar across the graft junction. Shown here is a grafting-based strategy for identification of mRNAs that move from shoot (scion) to root (stock), from genotype 2 to genotype 1, using a scion:stock=genotype 2:genotype 1 heterograft. The same strategy can be used to identify transcripts that move from shoot to root from genotype 1 to genotype 2 using a genotype 1:genotype 2 graft. Transcripts that move root to shoot can be identified by analysing mRNAs in shoot tissue. Natural grafts, such as those established between the parasitic dodder plant and its host plants, can be used in place of artificial grafts. A key challenge in all such approaches is how to assign transcripts to each genotype; methods for doing so are based on (1) SNP (single nucleotide polymorphism) identification or on (2) the alignment to different reference genomes. For grafts from the same species, or similar genotypes, SNPs can be used to distinguish between genotypes and thus identify the source genotype of each transcript (1). For grafts between different species, mapping (2) each RNA-Seq read to the genome assemblies can be an effective method for determining which transcripts are specific to one species.

In plants, cell-to-cell mRNA movement is facilitated by plasmodesmata (10, 25). On sub-cellular scales, movement of mRNAs occurs via diffusion and trafficking by motor proteins (26). However, for larger distances, these modes of transport become less effective; for instance, an mRNA travelling a distance of 1 cm by diffusion would require approximately 1000 days, approximately 20 days by non-selective transport along microtubules, and approximately 2 days by selective active transport, based on published values for these processes in neurons (26, 27). Taking the movement through plasmodesmata for inter-cellular transport into account would lead to longer travel times (28, 29). Long-distance transport of mRNAs likely occurs through the phloem (30–32); mass flow reduces the transport time from several days to minutes and comfortably within the expected half-lives of most mRNAs (33, 34). Several studies have reported hundreds of mRNAs in the phloem (14, 35–37). Thus, upon reconnection of their vascular system after grafting, we might expect hundreds of phloem-delivered mRNAs to be able to cross the graft junction, Figure 1. Indeed, studies over the last decade have identified hundreds to potentially thousands of graft-mobile mRNAs in a variety of plant species (1–7).

The ability of mRNAs to travel over long-distances coupled with reports of mRNAs acting non-cell autonomously has given rise to the suggestion that mobile mRNAs may act as systemic regulators or signals (38–43). However, reports place the estimated number of transported transcripts of any individual mobile mRNA at less than 0.1% of the local pool size for most transcripts (3, 5, 7). If transcription of an mRNA resulted in a pool size of 1000 transcripts then the expected stochastic fluctuations would be greater than *±* 30 transcripts (typically over-dispersed compared to a Poisson distribution (44, 45)). For a ratio of 1:500 for mobile over local transcript numbers (46), the expected number of imported mobile transcripts of the same mRNA would be only 2, i.e. the number of mobile transcripts would be only a fraction of the expected transcriptional noise (44) and several orders of magnitude less than the total pool size. Whilst the above values are derived from bulk data and may well differ for specific cell types, this nevertheless raises questions about how such signals can be recognised and processed (decoded). Ideas from information theory have been put forward as a means of analysing mRNA signaling as a communication system (17, 47).

For elucidating their signaling capacity and biological function, it is critical to know which mRNAs move, when they travel (under which conditions), and from where to where. Furthermore, determining which mRNAs are translocated might allow for features associated with mobility to be identified, such as physico-chemical properties or sequence and structural patterns (motifs, zip-codes). For instance, we previously identified a potential mobility motif that was enriched in a low abundance subset that turned out to relate to mRNA stability (48), in line with recent findings that suggest that mRNA localisation can be explained by mRNA stabilisation (26). Also, a tRNA-like structure (TLS) was identified as a transport motif and experimentally confirmed as being key for mobility (46, 49). There is therefore significant interest in the detection of mobile mRNAs and many reports and datasets now available (50). Data-mining such resources presents an exciting opportunity to identify features and mobility motifs that would shed light on the selectivity of transported mRNAs and greatly enhance our ability to engineer constructs for transgene-free genome editing (46). Nevertheless, extensive data-mining efforts, including our own, have failed to find predictive features for mobility in sequences of currently available datasets (51).

Here, we sought to establish a high-confidence dataset of mobile mRNAs, with the goal of using this dataset for learning the determinants of mobility to develop and train predictive models. However, looking in detail into the data supporting the annotation revealed several confounding factors in current methods for detecting mobile mRNAs. Our re-analysis of available datasets suggests that the reported variability between experiments (16) is potentially a consequence of the criteria that have been employed to classify mRNAs as being mobile. We have previously shown that criteria based on absolute read counts lead to a dependence on read-depth for simulated data (52) and we confirm this dependency for real data. We show that criteria based on mapping to genomes depend on genome assembly completeness and quality, and that other sources of variation, such as gene copy number, can bias the analysis. Therefore, whilst we and others have placed great care and effort in sample preparation (taking precautions to reduce contamination during RNA harvesting and sequencing (2, 5)) and data processing (using stringent quality filters, conservative thresholds and high-quality genome assemblies (2, 6)), armed with new insights into plant genomes and the availability of advanced data analysis tools, we must consider the possibility that previous studies may have over-estimated the number of mobile mRNAs from available data.

## Results

### SNP-based detection methods for mobile mRNAs may be confounded by technological noise

Identifying which transcripts belong to which genotype in grafted plants (between two different genotypes) requires a means of distinguishing them. Choosing closely related genotypes that differ in known positions between their genomes, single nucleotide polymorphisms (SNPs), allows for transcripts to be identified based on these allele-specific nucleotides in RNA-Seq data, Figure 1. Typically, a requirement is made for a defined number of RNA-Seq reads to have a SNP that corresponds to the alternative allele for a transcript to be assigned to a foreign genotype. These values are chosen carefully to reduce the risk of sequencing noise biasing the identification. Previously published criteria include at least one read if it is covering a minimum of two SNPs (3), or two (3), three (53) or four (2) reads covering a single SNP. When these criteria are met, the transcript is defined as mobile.

We asked how robust these criteria are to varying sequencing depths and different sources of noise. Sequencing has known technological limitations which lead to nucleotides occasionally being assigned to a different nucleotide, i.e. there are base-calling errors. Such occurrences are rare; Illumina sequencing machines produce base-calling errors at a rate of approximately 0.1 to 1% per base on average (54, 55). Sequencing providers often provide a quality assurance, for instance that 85% of the reads have a Phred quality score of at least Q30 (i.e. a base-calling error of less than 10^−3^). Thus, assuming an RNA-Seq experiment delivers 20 million reads using paired-end sequencing of 2 × 100 bp, we will have a total of 4 billion base calls of which we can expect up to approximately 4 million to be incorrect. Often, more stringent Phred score filtering is applied to enhance the quality, and reduce the number of errors (2). Base-calling errors are not the only potential source of change in the expected nucleotide identity. Prior to sequencing, reverse transcriptases can introduce base changes with an error rate of approximately 10^−5^ to 10^−4^; the reverse transcription reaction error can be different for different nucleotides, for instance a ‘G’ to ‘A’ bias (56, 57), and may exhibit a range of artifacts (58). In the above example, this could result in a further 400,000 potential nucleotide substitutions. Multiple effects give rise to differences between the sequenced fragments and the corresponding genome sequence. Some of these differences may appear in SNP positions and be indistinguishable from a SNP supporting the alternate allele. The alignment of RNA-Seq reads to the genomes of the two grafted plants is a process with two outcomes (endogenous genotype, foreign genotype), for which, with no further knowledge, the binomial distribution represents an optimal (maximum entropy, least-biased) probability assignment (59). We denote the probability of a SNP matching the alternate allele by *q*. Using a (cumulative) binomial distribution, we assessed existing mobile mRNA criteria by computing the probability of transcripts being classified as mobile based on sequencing noise, Figure 2.

**Fig. 2.**
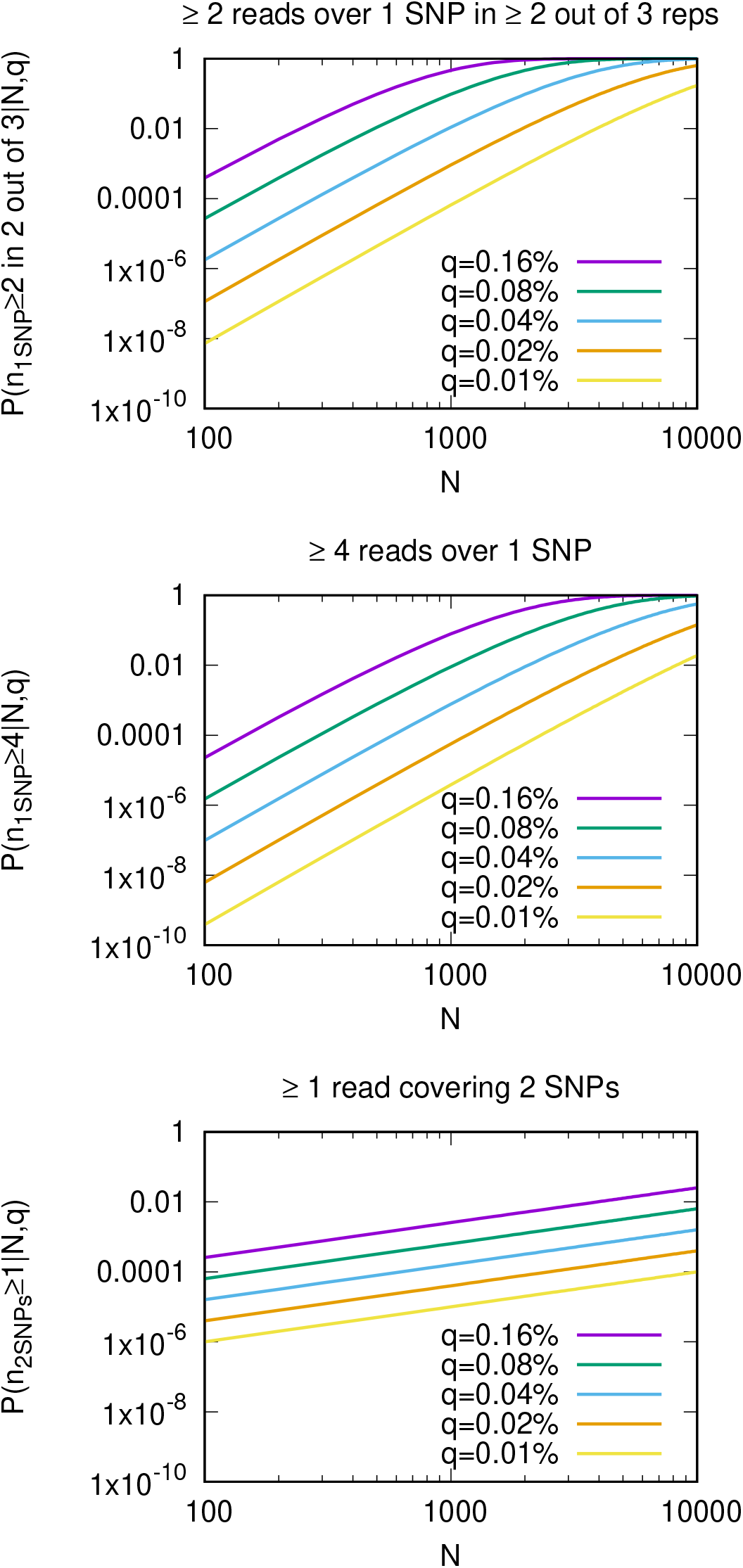
Published criteria for defining mobile mRNAs based on absolute read counts suffer from read-depth dependencies. These plots show the probabilities of transcripts being defined as mobile by chance. Three different mobile mRNA definitions (top, middle, bottom) and their dependence on read-depth (*N*) and on the rate of a SNP matching to the alternate allele (*q*) are depicted. The number of read counts over one SNP that correspond to the alternate allele is denoted by n_1SNP_, over two SNPs by n_2SNPs_. The probabilities were calculated using a cumulative binomial distribution, i.e. we account only for the nucleotides that correspond to the two alleles of interest. Note that both axes are on a log-scale. The requirement for co-occurring SNPs on one read is more stringent and less likely to occur by chance at higher read-depths. For low values of *q*, these criteria are robust up to moderately high (several hundred) read-depths and would be unlikely to occur by chance.

Criteria based on absolute read counts exhibit a read-depth dependency, Figure 2, resulting in a bias towards highly abundant transcripts being assigned as mobile (48, 52). We therefore investigated how many previously published mobile mRNAs identified by RNA-Seq are consistent with technological noise and read-depth. Table 1 lists how many mRNAs have been reported to be mobile and how many of these have occurrences of the alternate allele that are consistent with an assumed error rate. Two examples of error rates for the alternate allele, *q* = 0.1% and *q* = 0.01%, are provided. The hypotheses for whether the data can be explained by background errors or whether the experimental observations go beyond what would be expected from background errors alone (and are therefore potential candidates for mobile transcripts) were evaluated by computing the posterior probabilities and Bayes factors (52, 59, 60).

**Table 1.** Total numbers of reported mobile mRNAs in *Arabidopsis thaliana* (2) and *Vitis girdiana* (3) that can be explained by expected sequencing noise.

| | # of reported mobile mRNAs | # consistent with $q = 0.1\%$ | # consistent with $q = 0.01\%$ |
| --- | --- | --- | --- |
| *A. thaliana* (2) | 2006 | 1455 | 1086 |
| *V. girdiana* (3) | 1130 | 945 | 384 |

If the probability for the alternate allele, *q*, was purely a consequence of base-calling errors, we would expect *q* to be approximately a third of the base-calling error; for a Phred quality score cut-off of Q30 this would result in a base-calling error rate of 10^−3^ and a *q* of approximately 10^−3^*/*3 or 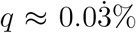. This *q* value falls within a range for which we find that a substantial number of previously identified mobile mRNAs (2, 3, 7) were classified based on numbers of RNA-Seq reads over SNP positions that are in line with what would be expected from sequencing noise, Table 1. We point out that we used the same value of *q* for each base in this analysis and did not take position-specific error rates into account. Nevertheless, this analysis suggests that, based on available RNA-Seq data, a large proportion of the annotated mobile mRNAs may lack statistical support.

### SNP-based detection methods for mobile mRNAs may be confounded by biological noise

If SNPs are located closely together then a single RNA-Seq read may cover more than one SNP (we will refer to these as co-occurring SNPs). Requiring RNA-Seq reads over co-occurring SNPs that support the alternate allele results in a less pronounced read-depth dependence than single SNP criteria, Figure 2. We therefore investigated RNA-Seq reads that covered multiple SNPs. The examination of reads over co-occuring SNPs revealed another confounding factor in the identification of mobile mRNA; several loci showed apparent heterozygosity in the homograft data, Supplemental Figure S1 and Supplemental Figure S2 and S3. While the *Arabidopsis thaliana* ecotype *Columbia* (Col-0) assembly is of excellent quality, the genome assemblies of different Arabidopsis ecotypes are not all of the same quality. For instance, a recent study of the *Landsberg erecta* (Ler-1) ecotype (61) of *Arabidopsis thaliana* identified 105 single copy genes present only in Ler-1 but not in Col-0, and 334 single copy orthologs that had an additional copy in only one of the genome assemblies. It has been estimated that 10% of the annotated genes in Arabidopsis have copy-number variation (61, 62). Differences in gene copy numbers can lead to reads not mapping correctly, which gives rise to pseudo-SNPs and pseudo-heterozygosity (61, 62). Whilst challenging to quantify, a high fraction of reads with variation at specific positions within a single genome can be indicative of pseudo-heterozygosity (61, 62). Looking in further detail on a case-by-case basis, we found several mobile mRNAs assignments that are possibly caused by this pseudo-heterozygosity that we identified in the homograft data, Supplemental Figures S1, S4. Puzzlingly, some experimental samples seem to exhibit pseudo-heterozygosity in only one of the tissues (root or shoot tissue). This is possibly a consequence of gene duplicates (paralogs) having differential expression in different tissues. Thus, in addition to technological noise, there are also biological causes that could be falsely interpreted as SNPs of an alternate allele.

### Technological and biological noise account for a majority of previously identified mobile transcripts

Having identified sources of noise in addition to base-calling errors, we sought to estimate the frequency with which a SNP would be identified as matching to the alternate allele when the alternate allele is not present, i.e. the background noise level. This value can be estimated from homograft data. From available Arabidopsis homograft data (2) we inferred a background rate for the alternate allele of around 0.1%. The estimated rate for the alternate allele of *q* = 0.1% is higher than would be expected from base-calling errors for a Phred score threshold of Q30 and suggests that other effects, such as reverse transcriptase artifacts (58) or bioinformatic ambiguities (arising from recently duplicated genes, conserved or repetitive regions in large gene families, gene copy number variation, genome completeness and genome quality) cannot be neglected. With a value of *q* = 0.1% we find that between 72% and 83% of annotated mobile mRNAs would not be distinguishable from expected errors in homograft data, Table 1. Supplemental Figure S5 shows how the uncertainty in the inferred error rate depends on the amount of data. We capture this uncertainty through probability distributions to inform inferences drawn from the data (52). Using SNP-specific distributions, derived from counts per SNP in homograft data (52), the statistical differences between heterograft and homograft decrease further.

To investigate the differences between SNPs and non-SNPs, we computed the nucleotide distributions at different positions. Given the expected low numbers of mobile mRNAs relative to local transcripts, most SNPs in mobile mRNAs will not have sufficient coverage to detect foreign alleles, Supplemental Figure S6. Those SNPs that have reads matching the alternate allele may provide evidence in support of the associated mRNA being from another genotype and potentially being trafficked into the sampled tissue. Figure 3 shows the distribution of reads that match to the alternate allele, *n*, over the total number of reads, *N*, for each SNP in the mobile population of two example datasets (2, 3). We can see that several SNPs have support for the alternate allele. We would expect these distributions to be different from other, non-SNP, positions in the sequence. However, looking at all neighbouring positions of SNPs in the mobile population and computing the number of reads with second most frequent nucleotide, *m*, over the most frequent and second most frequent nucleotides, *M*, we find little difference between SNPs and non-SNPs. More reads with nucleotides from the alternate allele at a SNP position should lead to a shift in the distribution towards higher *n/N* values, i.e. *n/N > m/M*, but this is not observed. This is in line with the above analysis that suggests that many reads that have been interpreted as supporting the alternate allele are actually consistent with sequencing noise.

**Fig. 3.**
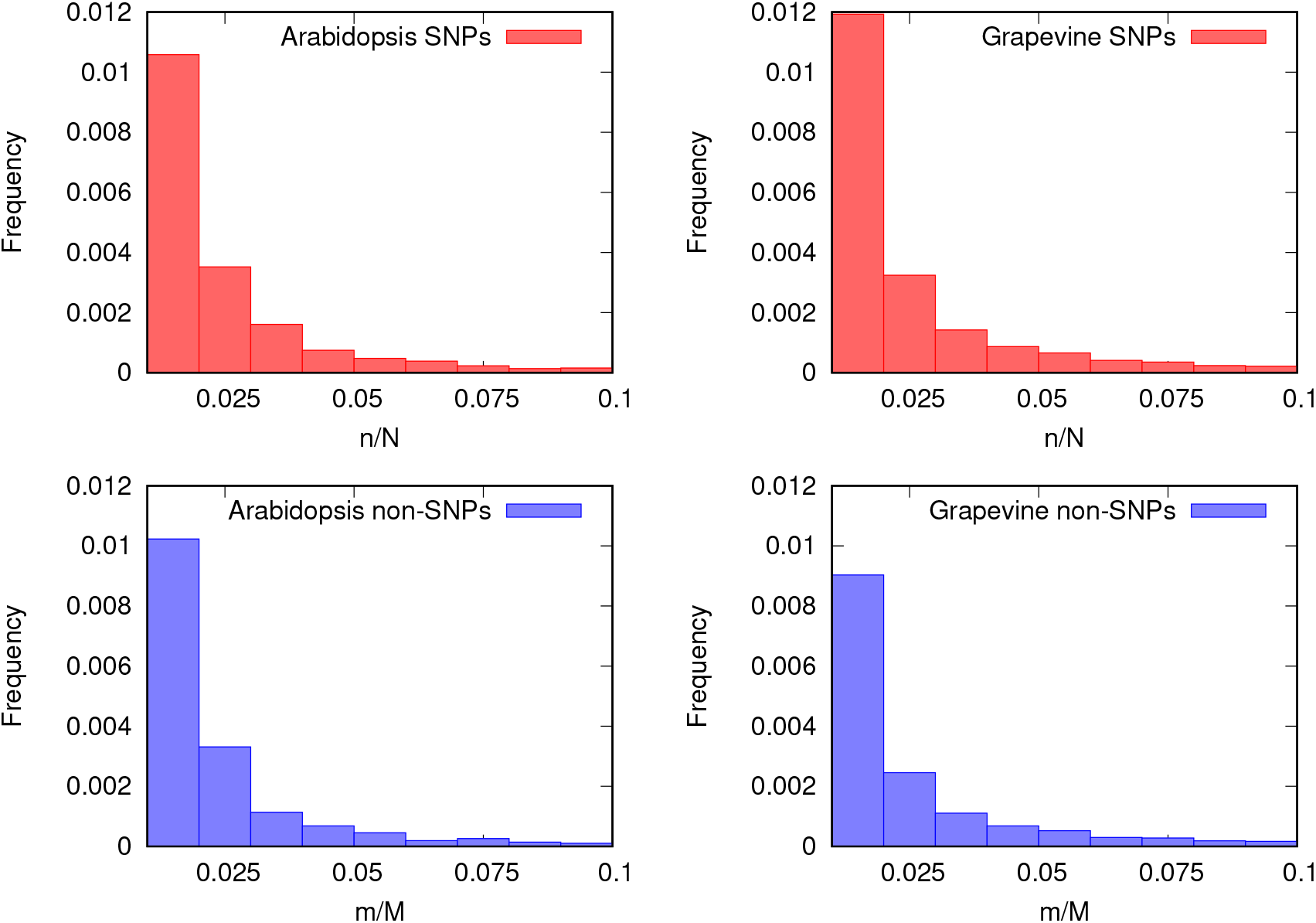
The distributions of nucleotides at SNP and non-SNP positions can be informative for evaluating the evidence for the alternate allele. Top (red): Histograms of the ratio of the number reads that match to the alternate allele, *n*, over the number of reads of local and foreign reads, *N*, for each SNP position in the mobile population on examples from Arabidopsis (2) and grapevine (3) on the left and right, respectively. Several SNPs have reads that match to the alternate allele. The full distributions are depicted in Supplemental Figure S6. Bottom (blue): Histograms of the ratio of the number reads that match to the second most frequent nucleotide, *m*, over the sum of the number of reads over the most frequent and second most frequent nucleotide, *M*, for neighbouring positions to SNPs. An overlay of the distributions is given in Supplemental Figure S7. In these examples, the SNP distributions are similar to non-SNP distributions, suggesting that both may be a consequence of the same technological noise.

To summarise, we find that technological sequencing noise and biological variation is challenging to discern from evidence for SNPs from alternate alleles. Basing the identification of mobile mRNAs on absolute numbers of RNA-Seq reads covering a particular SNP that support a specific genotype could result in technological noise from sequencing machines and genome differences being mistakenly interpreted as evidence for mobile mRNAs. These ideas are summarised in Figure 4. Identifying reads with multiple SNPs can provide greater confidence in the read being from a different genotype.

**Fig. 4.**
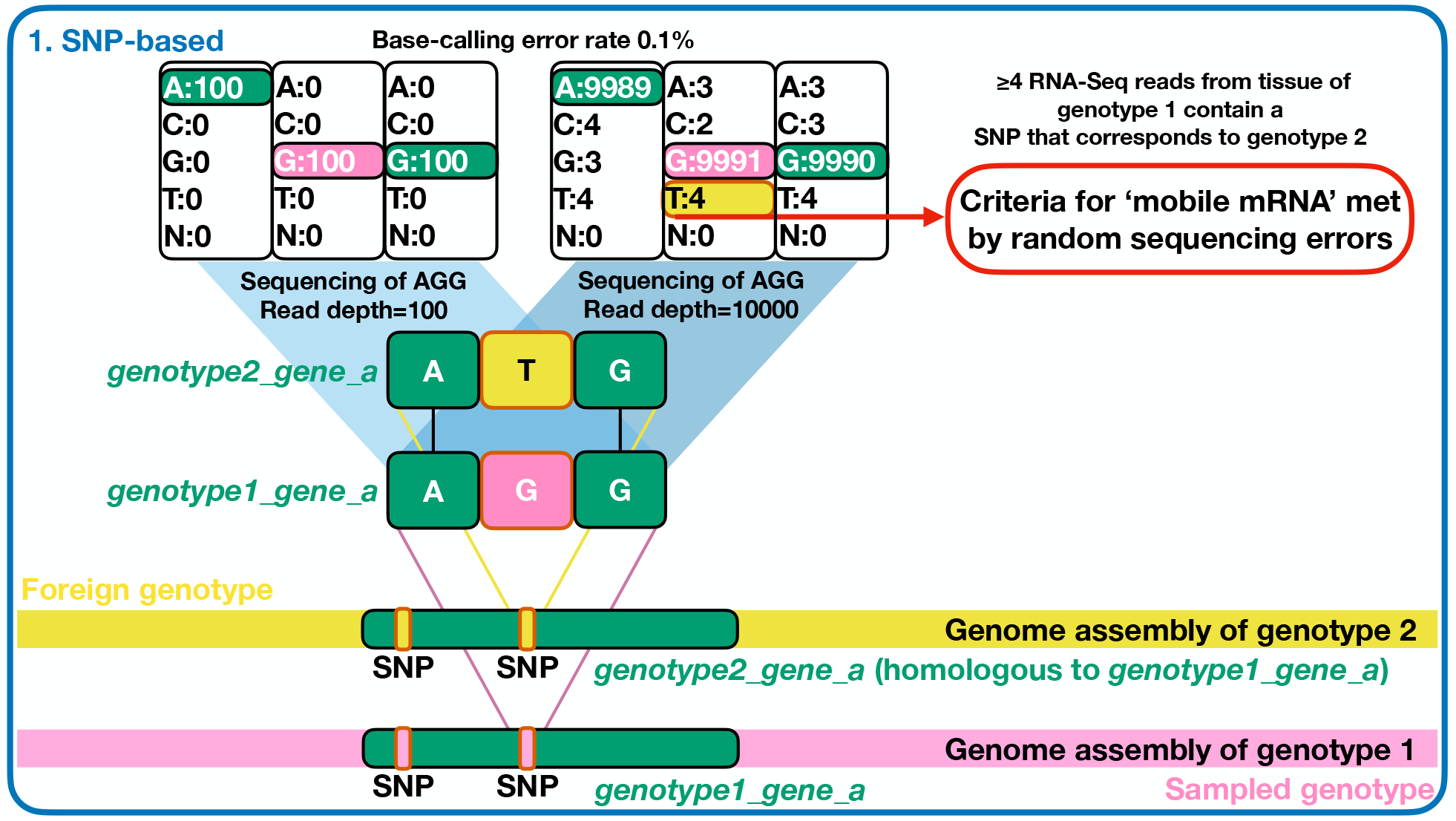
Technological noise can lead to challenges in the assignment of RNA-Seq reads. In SNP-based methods of mobile mRNA detection it is important to be able to differentiate between sequencing-associated errors and genuine SNPs. In the above case, an RNA-Seq read with a ‘T’ at the SNP position would be indicative of the read having come from the alternate allele, genotype 2. However, every position has an error rate and the higher the read-depth the more incorrect base calls are to be expected. Base changes could arise for reverse transcriptase or amplification steps, although their error rate is typically orders of magnitude lower than sequencing errors. Conserved regions in gene families can give rise to similar challenges in distinguishing mapping ambiguities from genuine SNPs. Defining an mRNA as being mobile based on thresholds of RNA-Seq reads that contain a SNP can result in base-calling errors and mapping ambiguities biasing the interpretation. To reduce the risk of such events occurring, further stringent filters can be applied (for instance using only SNPs that are bi-allelic (2)) or applying rigorous statistical comparisons (for instance estimating the allele calling frequencies and comparing them between homograft and heterograft (52)).

### Homologous sequences and the quality of genome assemblies can influence the identification of mobile mRNAs

As copy number variation and pseudo-heterozygosity may affect the evidence for alternate alleles in intra-species, SNP-based approaches (61, 62), we asked whether cross-species genome mapping approaches that do not rely on SNPs may have similar shortcomings with respect to genome quality and completeness. The quality of genome assemblies varies depending on sequencing and assembly strategies, as well as the underlying complexity of the genome (63). In Table 2 we list some genome completeness estimates for the genome versions that were used in previous mobile mRNA studies. Note that these genomes are constantly being improved and newer versions have been published, and that the completeness in gene-rich regions is what is important for aligning transcripts, not the overall completeness.

**Table 2.** Completeness of genome assemblies used in mobile mRNA studies.

|  | Genome size | Assembly size | %Assembled | Year of Publication |
| --- | --- | --- | --- | --- |
| *A. thaliana* Col-0, TAIR10 (64) | 135 Mb | 119.5 Mb | 88.5% | 2012 |
| *A. thaliana* Ler (61) | 135 Mb | 117 Mb | 86.7% | 2016 |
| *N. benthamiana* (65) | 3 Gb | 2.412 Gb | 80.4% | 2012 |
| *S. lycopersicum* (66) | 900 Mb | 760 Mb | 84.4% | 2012 |
| *V. vinifera* (67) | 500 Mb | 473.8 Mb | 94.8% | 2007 |

A typical mapping-based bioinformatics pipeline for analysing between-species grafts starts by mapping the reads to the reference genome of the sampled tissue (genotype 1 in Figure 1). Any unmapped reads can then be mapped to the reference genome of the potential source tissue (genotype 2 in Figure 1). We built a simple model, Supplemental Figure S8, of this process in which the probability of a RNA-Seq fragment mapping to a genome depends on the completeness (in terms of genes) of the genome assembly. Under this assumption and incomplete genome assemblies, it is possible that an RNA-Seq read from one genome is falsely assigned to the other genome, and thus considered to be mobile. In such cases, additional steps can be taken to identify reads potentially mapping to unassembled parts of the genome, such as aligning to transcriptome data.

We investigated the pipeline used to identify transcripts that travel from a *Nicotiana benthamiana* scion to a *Solanum lycopersicum* (tomato) rootstock (6). At that time, the tomato genome had approximately 15% of the genome unassembled, Table 2. Stringent criteria were therefore used to analyse the data to mitigate effects of using an incomplete assembly. To identify potential *N. benthamiana* transcripts in tomato tissue, first all RNA-seq reads from tomato stock samples matching the tomato reference genome were excluded. To account for the incomplete tomato reference genome, remaining reads were then matched to an RNA-Seq transcriptome generated from an ungrafted tomato sample (6). In this step, any reads from the grafted tomato stock that matched exactly to reads from the ungrafted tomato sample were also excluded. Having filtered all reads based on existing knowledge of the tomato genome and transcriptome, remaining reads were mapped to the *N. benthamiana* genome. Reads mapping to *N. benthamiana* were classified as having originated from mobile mRNAs. However, we found that many of these reads that did not map to the tomato genome all aligned to small regions of the *N. benthamiana* genome, and that coverage was highly uneven over exons, Supplemental Figures S9 and S10. Furthermore, blasting the reads identified as being from *N. benthamiana* against the whole of NCBI nucleotide database found 100% matches to highly-conserved sequences contained within many genomes, including *N. benthamiana* and Solanaceae species, in particular to 18S ribosomal RNA genes. It is possible that the criteria chosen for comparing RNA-Seq reads between grafted and ungrafted tomato were overly stringent, thus failing to exclude reads from the heterograft dataset that may have actually been from tomato. Such highly-conserved sequences from tomato could match to the *N. benthamiana* genome and be incorrectly identified as mobile. Coverage over the full length of the mRNA may help reduce the risk of reads mapping to isolated regions being potentially misinterpreted, Supplemental Figure S9. As for SNP-based approaches, it is possible that read-depth can bias the interpretation of RNA-Seq data from grafts between different species, Supplemental Figure S11. For instance, approximately 30% of the *Arabidopsis thaliana* transcriptome has been reported to move into *Cuscuta pentagona*, while only 9% of the tomato transcriptome moves to Cuscuta (1). However, there is a large difference in the amount of transcriptome sequencing data for tomato (6 Mb) and Arabidopsis experiments (2 Gb), and it is known that different coverages lead to different detection level (68–70).

In summary, we find that the completeness of genome assemblies and the presence of highly-conserved regions can introduce significant challenges for the identification of mobile mRNAs.

## Discussion

In plants, local cell-to-cell transport of mRNA occurs via plasmodesmata (10, 25), whereas long-distance translocation is thought to be facilitated by the phloem (15, 19, 30–32). The transport of several mRNAs, both cell-to-cell and over long distances, has been experimentally validated by methods that include the detection of the mobile mRNA directly, a translated protein, or a phenotypic change distal to the site of transcription of either an endogenous or transgene-derived sequence. The evidence for transport and translation of selected mRNAs is compelling (19, 46, 49) and the movement of mRNAs has been experimentally confirmed (5, 6, 10, 42). Based on RNA-Seq studies, over a hundred to several thousands of transcripts have been reported to be transported over long-distances in plants (1–7).

Here, we re-analysed RNA-Seq data and reviewed associated bioinformatic pipelines that have been used to identify mobile mRNAs. Our findings suggest that the evidence from RNA-seq data supporting many candidate mobile mRNAs cannot be discerned from technological noise, biological variation and genome assembly issues. This is most simply illustrated by comparing statistical properties of homograft and heterograft data or SNPs and non-SNPs, Figure 3. Although several mRNAs that have been identified as being mobile based on RNA-Seq data have been experimentally validated (2, 5, 46, 71), we cannot rule out that others might be false positives.

If current mobile mRNA detection methods based on short-read RNA-Seq are indeed susceptible to technical and biological noise, we would expect a pronounced read-depth dependence for SNP-based approaches and considerable variation between experiments due to the nature of sequencing associated errors. This read-depth dependence is likely to be the underlying cause for the observed dependence on abundance and apparent non-selective movement (48), i.e. abundance is not causative for mobility *per se* but is rather a consequence of how mobile mRNAs had been defined. These ideas are summarised in Figure 4. Interestingly, in mammalian systems, mRNA transport was also found to be pervasive across the genome, in a non-selective and expression dependent manner (72).

We conclude that identifying putative mobile mRNAs from grafted plants and RNA-Seq is far from trivial. The presented analysis identifies that there are significant computational challenges and technical hurdles facing researchers trying to identify alternate alleles associated mRNAs from RNA-Seq data. These concepts are depicted in Figures 4 and 5. Once a transcript has successfully been identified from scion samples as originating from the stock tissue, or *vice versa*, then a common hypothesis is that it may have been transported. This is one possibility, however, there are others; even after stringent controls, the possibility of low levels of contamination can rarely be excluded. The current criteria used to define a transcript as being mobile should therefore rather be viewed as criteria for a transcript being from a foreign genotype. However, as demonstrated, such criteria lead to read-depth dependencies, Figure 2, and are best avoided. Using short-read RNA-Seq to identify mobile mRNAs has limitations, particularly when the transported numbers are below the typical noise levels of this technology. Base-calling errors, reverse transcriptase errors, differences in coverage, batch effects, potential contamination, copy number variation and genome reference quality all represent alternate hypotheses that also consistent with the available data. It thus becomes important to evaluate different hypotheses to explain the RNA-Seq data rather than to define transcripts as mobile based on chosen criteria. The presented work describes the complexities in identifying mobile mRNAs, thus raising questions about the extent of mRNA signaling and highlighting that we still have a lot to understand about long-distance mRNA transport and function in plants.

**Fig. 5.**
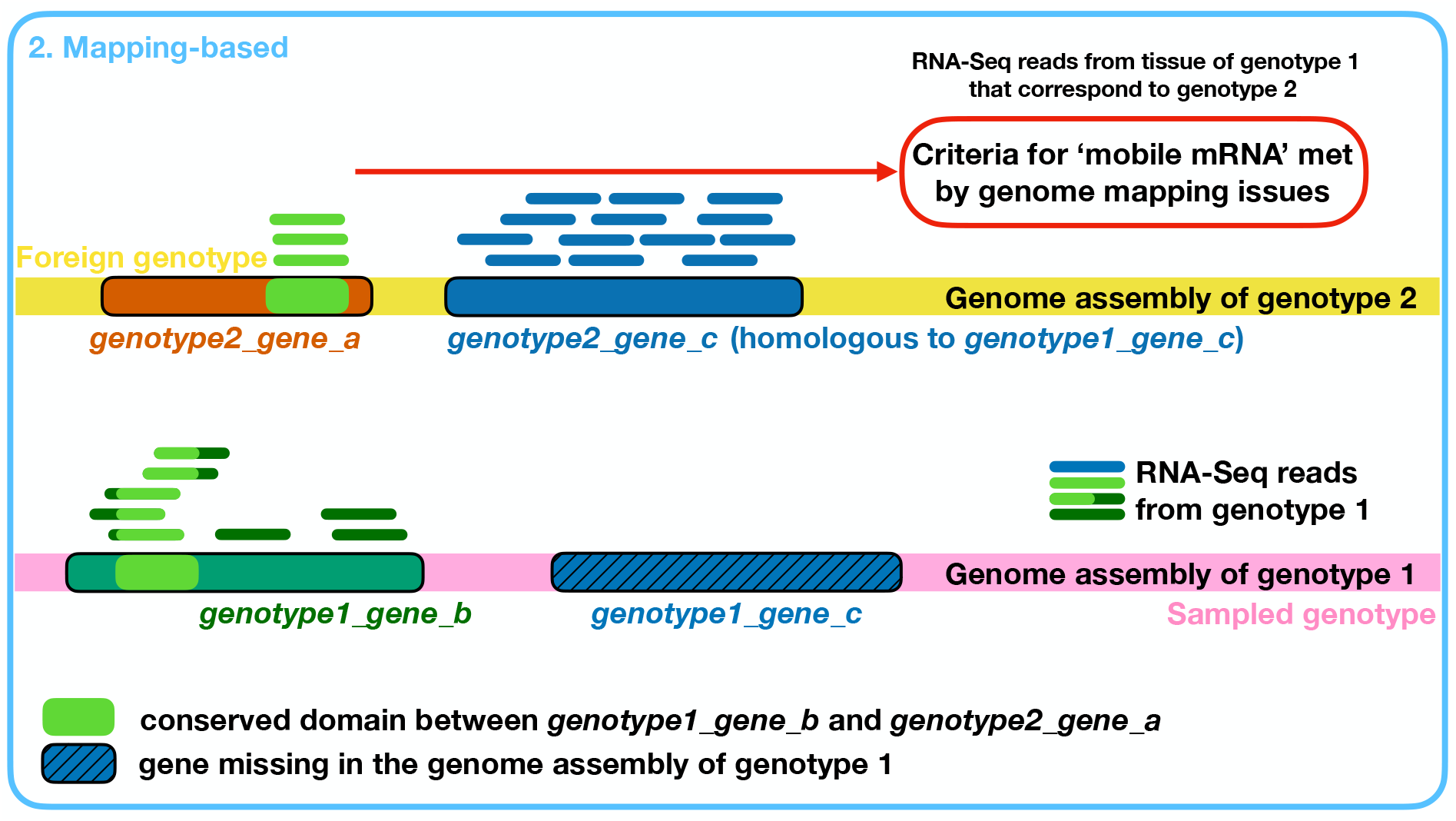
Genome complexity and genome quality can lead to mapping challenges. Orthologous sequences (light green) can result in some RNA-Seq reads aligning to a different gene and different genotype. Genome assemblies that are not complete (telomere to telomere) from exactly the same genotype as used for grafting can result in potential mis-mappings. The shaded blue gene in genotype 1 is missing in the reference genome assembly, resulting in RNA-Seq reads from this transcript being mapped to genotype 2.

## Methods

In this study we used the following published datasets, including the archived reads from NCBI as follows. *Cuscuta pentagona* (1): PRJNA257158. This dataset was incomplete and partly corrupt. *Vitis vinifera* (3): SRP058158 and SRP058157. *Solanum lycopersicum, Nicotiana benthamiana* (6): SRP111187. *Arabidopsis thaliana* (2): PRJNA271927. *Citrullus lanatus* L. (4): PRJNA553072. We also used the deposited Supplementary datasets to obtain the numbers of identified mRNAs. For each of the graft studies, we downloaded the reference genome sequence that matched the one that was used in the original paper with the same annotations; these were all publicly available in Ensembl plants (73). The raw reads were mapped to the references using hisat2 (v.2.1.0) (74), and processed by samtools (v1.9) (75), the expression levels were quantified with Stringtie (v1.3.5) (76). The variants were called with bcftools (v1.10.2) (75) using ‘bcftools mpileup -A -q 0 -Q 0 -B -d 500000 –annotate FORMAT/AD, FORMAT/ADF, FORMAT/ADR, FORMAT/DP, FORMAT/SP, INFO/AD, INFO/ADF, INFO/ADR’. The NCBI nucleotide database was downloaded on (21/Oct/2022) and (blast+ v2.9.0) (77) was for alignments. Probabilities of errors were calculated from a standard binomial distribution. The statistical comparison of error rates was performed using baymobil (52, 60).

## Acknowledgements

We are grateful to Prof Staiger (Bielefeld University), Dr Wilfried Haerty (Earlham Institute), Prof Korbinian Schneeberger (LMU Munich), Dr Melinda Mayer (QIB), Prof Zagrovic and Dr Polyansky (Max Perutz Labs, Vienna) for discussions, insightful comments and constructive feedback on previous versions of the manuscript. The presented re-analysis and the insights derived from it would thus not have been possible without the availability of raw data and we thank all authors who deposited their data, meta-data, and methods in public repositories.

PP: Investigation, Formal analysis, Writing - Original Draft, Writing - Review & Editing, Conceptualization, Visualization. MT: Investigation, Software, Formal analysis, Methodology, Writing - Review & Editing, Conceptualization. FH: Software, Methodology, Writing - Review & Editing, Conceptualization, Visualization. RV: Methodology, Writing - Review & Editing. MH: Writing - Review & Editing. HT: Writing - Review & Editing. FA: Writing - Review & Editing. ES: Writing - Review & Editing. SG: Writing - Review & Editing. MF: Writing - Review & Editing, Supervision. DW: Writing - Review & Editing. CF: Conceptualization, Writing - Review & Editing. JK: Conceptualization, Writing - Review & Editing, Funding acquisition. FK: Conceptualization, Writing - Review & Editing, Supervision, Funding acquisition. RM: Conceptualization, Formal analysis, Visualization, Writing - Original Draft, Writing - Review & Editing, Supervision, Funding acquisition

No competing interests.

