## Supplemental Figures for "Re-analysis of mobile mRNA datasets highlights challenges in the detection of mobile transcripts from short-read RNA-Seq data"

### Supplementary materials

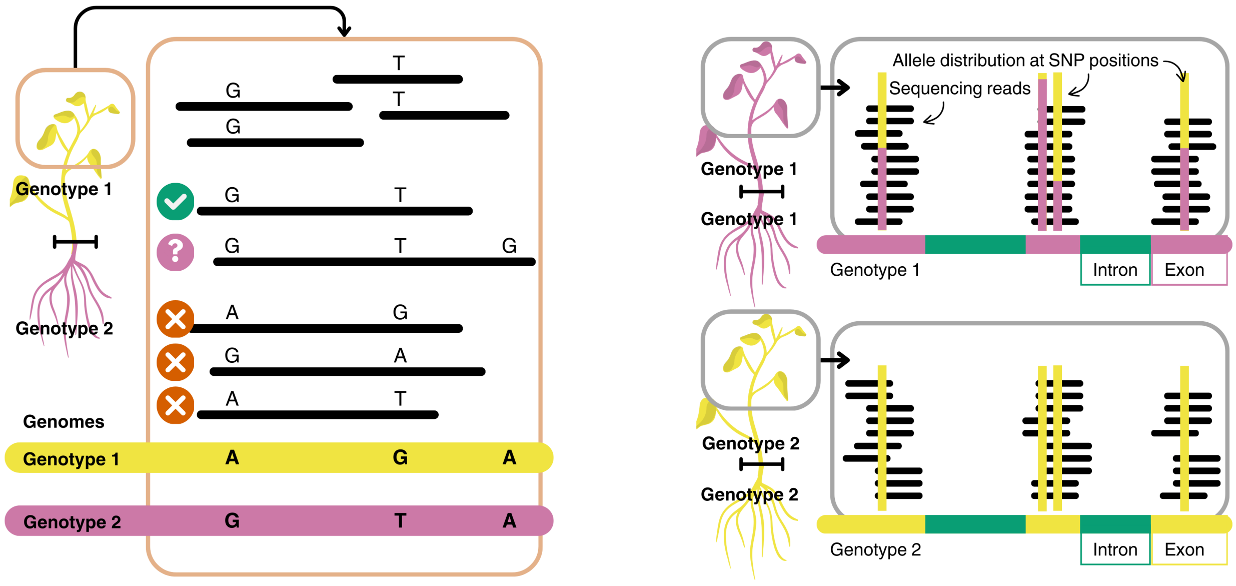


(a) (b)

**Supplemental Fig. S1.** Allelic differences in multiple SNPs per read and the appearance of heterozygosity (in homozygous species) can be used to check the viability of SNPs and exclude potentially problematic transcripts from the analysis. (a) SNPs can be in close proximity, and therefore it can happen that several SNPs are recorded in the same RNA-Seq read. In this example, genotype 1 has three SNPs very close to each other: A, G and A (yellow bar). In genotype 2, we find G, T, A (magenta bar) in those positions. In this schematic example, reads from the shoot of Genotype 1 are mapped to Genotype 2. If all covered loci carry the allele of Genotype 2, we are observing evidence for the read being from Genotype 2 and the associated transcript being potentially mobile. On the other hand, if only one loci carries the allele of Genotype 2, the outcome is inconclusive, as it may indicate sequencing errors. (b) $A.\ thaliana$ is a selfing species, so we expect homozygocity at all positions for all reads mapping to the genome at all positions. However, for duplicated genes (magenta) in Genotype 1, which may be single copy genes in Genotype 2 (yellow), short read sequencing and mapping to Genotype 2, can give rise to what appears to be heterozygocity. When there are two alleles present in the homograft data (magenta and yellow), we may be observing pseudo-heterozygocity. See also Supplemental Figure S2.

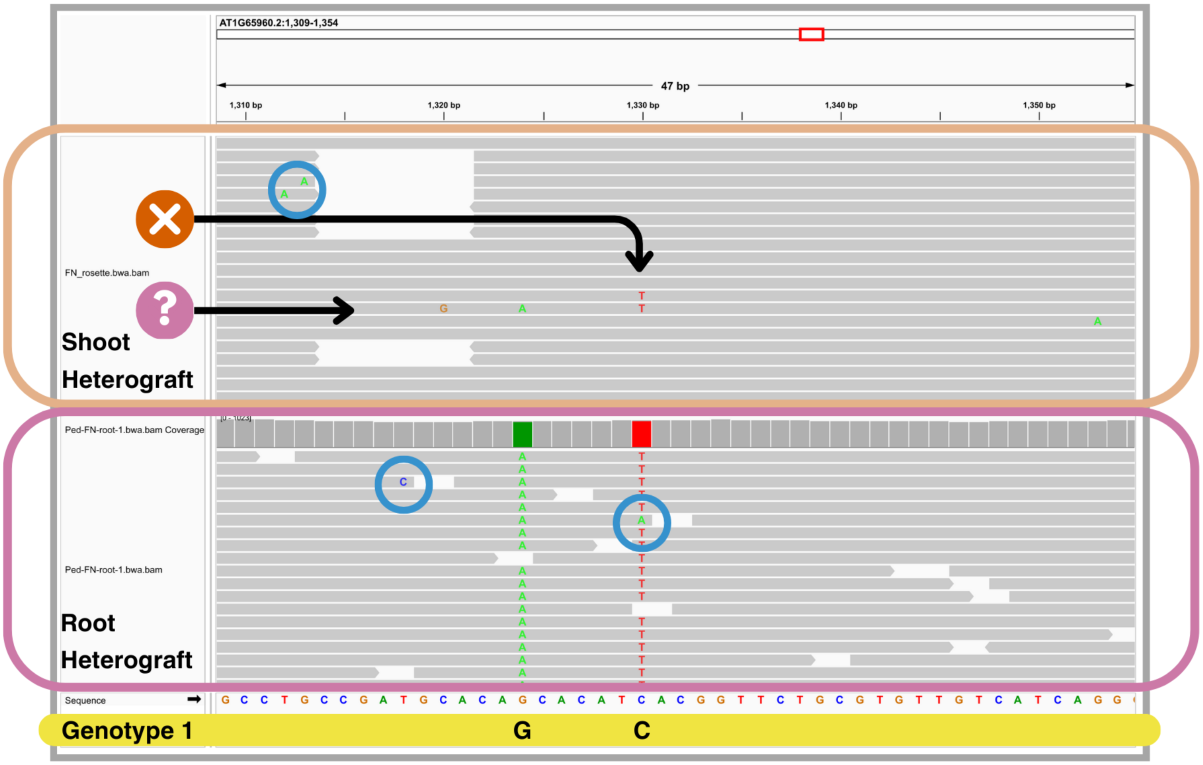


**Supplemental Fig. S2.** An example of the co-occurring SNPs. There are two SNPs, G and C in Genotype 1 (Col) and A and T in Genotype 2 (Ped). There is a likely two sequencing errors in the root sample, a C at a non-SNP position and an A at a SNP position (both highlighted in blue circles). In the shoot sample we see evidence for mobility at the SNP level but in one case the second SNP is not present and in the other case another sequencing error has occurred (G). Three further sequencing errors (two As on the top left, one A on the right) are also present in the shoot. See Figure 1 for further explanations. This is an annotated screenshot taken in IGV (78). Data taken from (2).

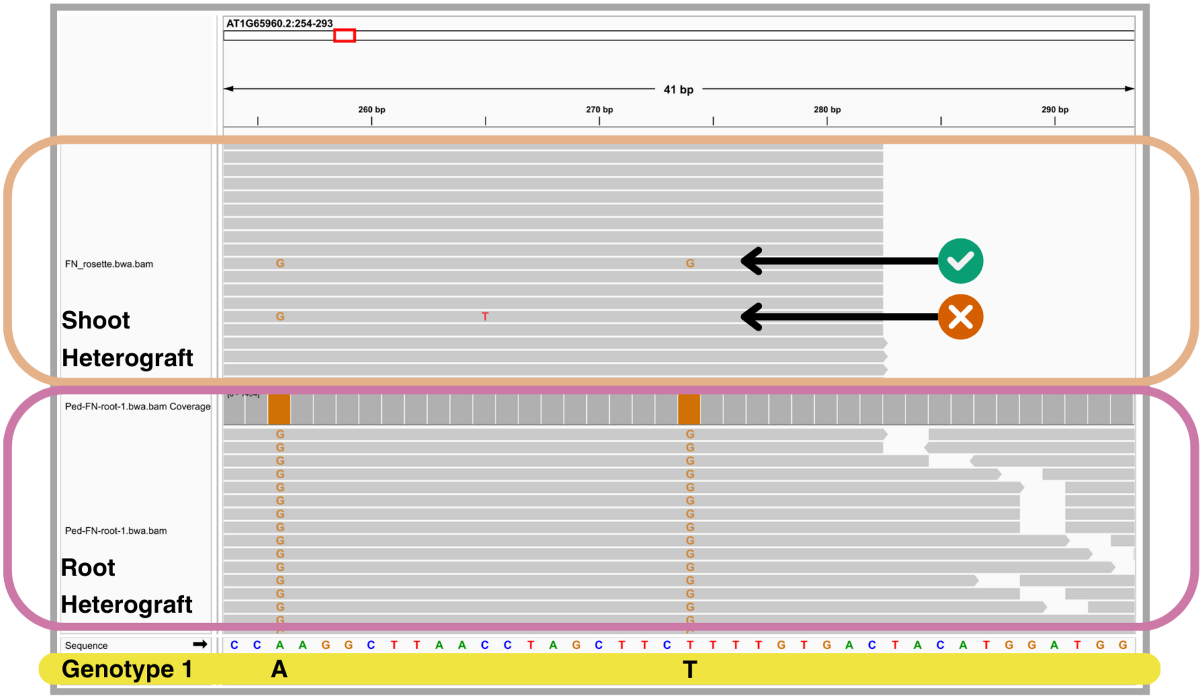


**Supplemental Fig. S3.** An example of the consistent and inconsistent co-occurring SNPs. See Supplemental Figure S1. This example shows two positions, A and T in Genotype 1 (Col) and G and G in Genotype 2 (Ped), for which some reads support the alternate allele (green tick), whereas others are likely sequencing errors (red cross). In the latter case, one G is in the correct position but the other G is not present and a further mismatch (T) has occurred. This is an annotated screenshot taken in IGV (78). Data taken from (2).

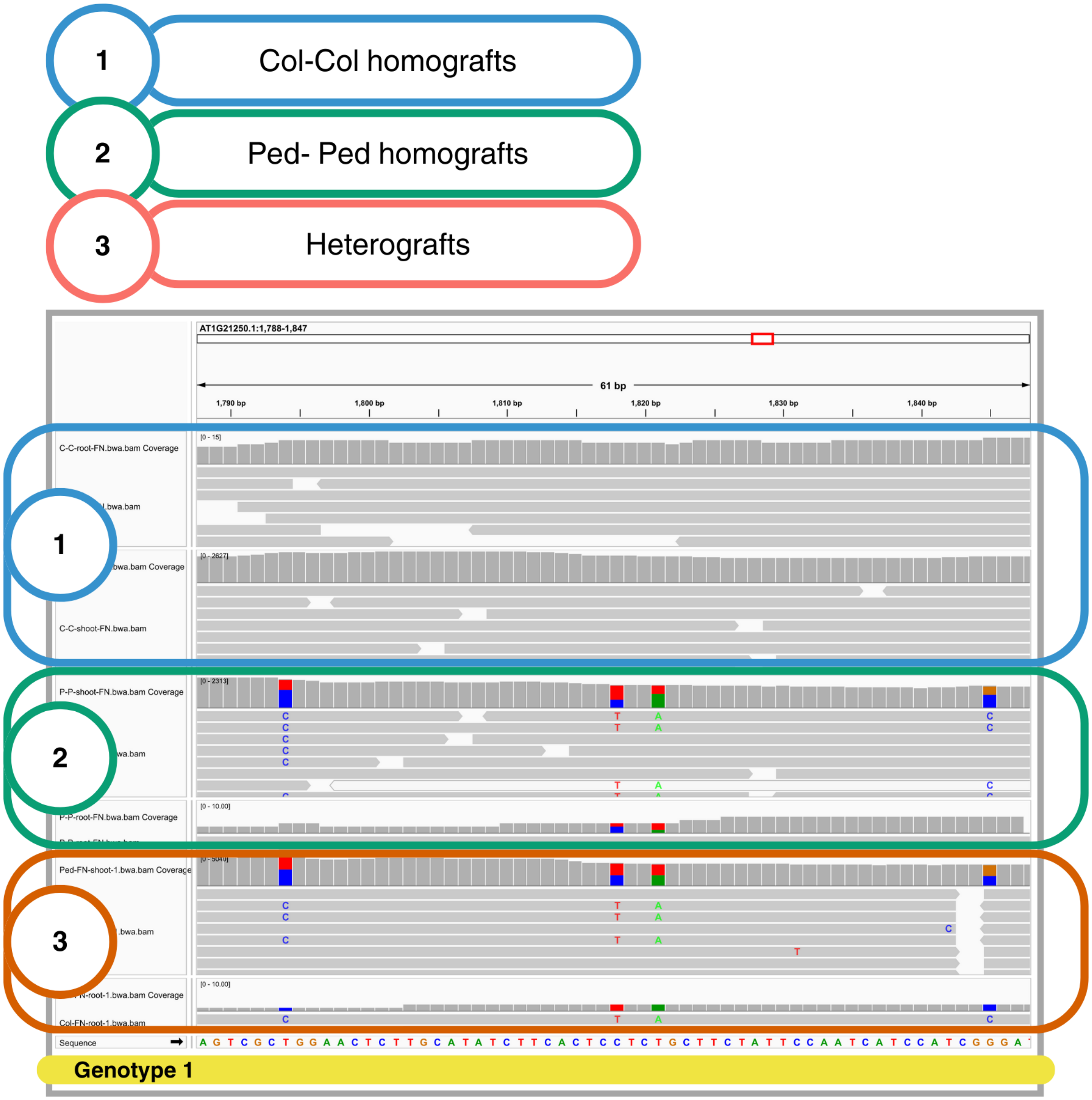


**Supplemental Fig. S4.** There are a number of genes in Ped that show apparent heterozygocity, both in the homograft (2) and heterograft (3) datasets. At the highlighted positions there are distinct populations of alleles (depicted as red/blue and red/green bars). This is possibly due to the gene being duplicated in Ped, resulting in pseudoheterozygocity.

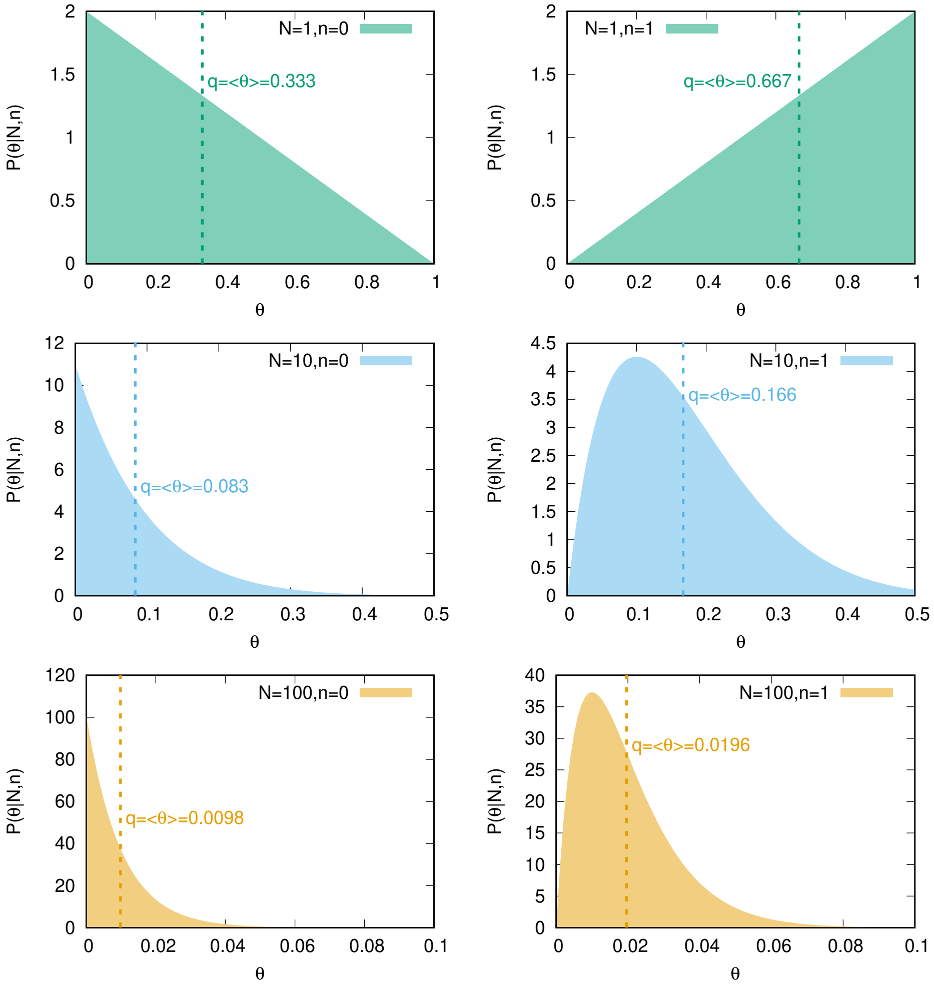


**Supplemental Fig. S5.** The uncertainty in the determination of the error rate depends on the read depth. $N$ is the total number of RNA-Seq reads over a SNP and $n$ is the number of RNA-Seq reads with a nucleotide matching the alternate allele at this SNP position. The plots show the posterior probability distribution over the inferred error rate, $\theta$, starting from a uniform prior (52). The most likely values of $\theta$ are consistent with $n/N$, whereas the expectation value over $\theta$, $q = \langle\theta\rangle$, represents the best estimate of the error from the data. An empirical approximation for ranking mobile mRNAs based on such considerations was previously derived using a binomial distribution (2). This equation correctly captures the key dependencies (2, 52). We recently extended this approach and presented an exact solution that takes the full uncertainty (the full distributions as shown above) into account (52).

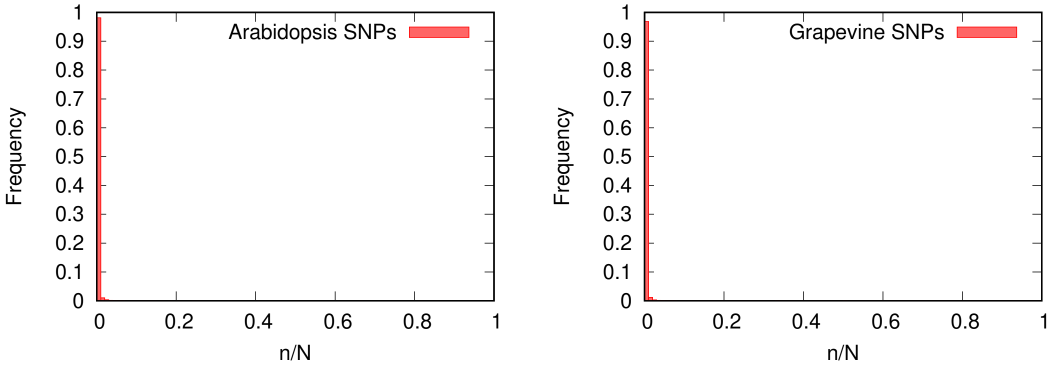


**Supplemental Fig. S6.** Most SNPs in mobile mRNAs do not have evidence for the alternate allele. Histograms of the ratio of the number of reads that match to the alternate allele, $n$, over the number of reads of local and foreign reads, $N$, for each SNP position in the mobile population on examples from Arabidopsis (2) and grapevine (3).

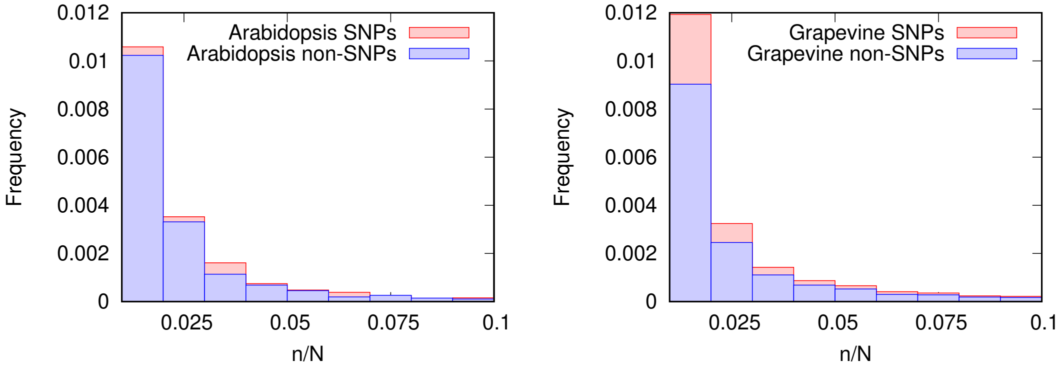


**Supplemental Fig. S7.** Overlay of the ratios of nucleotides at SNP and non-SNP position. Histograms of the ratio of the number reads that match to the alternate allele, $n$, over the number of reads of local and foreign reads, $N$, for each SNP position in the mobile population on examples from Arabidopsis (2) and grapevine (3) on the left and right, respectively. For non-SNP positions the ratio of the number reads that match to the second most frequent nucleotide, $m$, over the sum of the number of reads over the most frequent and second most frequent nucleotide, $M$, is depicted. The frequencies are somewhat different but the actual values of $n/N$ are similar for SNP and non-SNP positions.

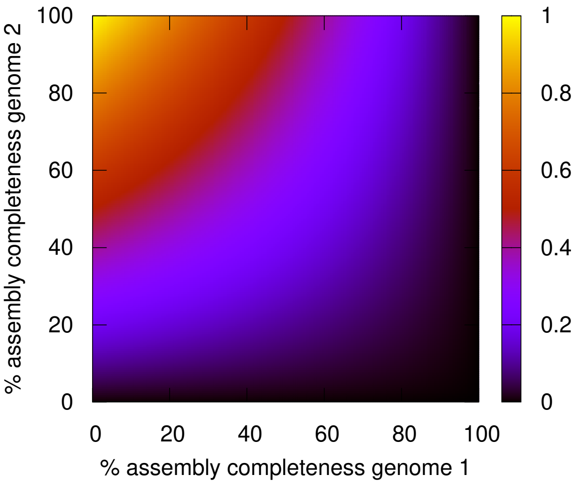


**Supplemental Fig. S 8.** Genome assembly incompleteness can lead to mapping challenges that may result in the identification of non-mobile transcripts being defined as mobile. We assume a simplified procedure in which RNA-Seq reads are first mapped to genome 1 and those that don not align to genome 1 are then aligned to genome 2. If the RNA-Seq reads match to regions of genome 2, the corresponding transcript is considered to be mobile. If the genome assembly of genome 1 was complete, then there would be no reads left to align to genome 2. If the assembly of genome 2 had low completeness, few RNA-Seq reads would be successfully aligned to it. In this simple model, the probability of a RNA-Seq fragment mapping to genome 2 depends on not mapping to genome 1 (denoted by a bar over 1), $P(\bar{1}) = (1 - c_1)$, and the probability of mapping to genome 2, $P(2) = c_2$, where $c_1$ and $c_2$ are the completeness values for the genome assemblies. $P(\mathrm{mobile}) = P_{\mathrm{max}} P(\bar{1}) P(2)$. The variable term $P(\bar{1})P(2)$ is shown as a heatmap as a function of the completeness of genome 1, $c_1$, and genome 2, $c_2$.

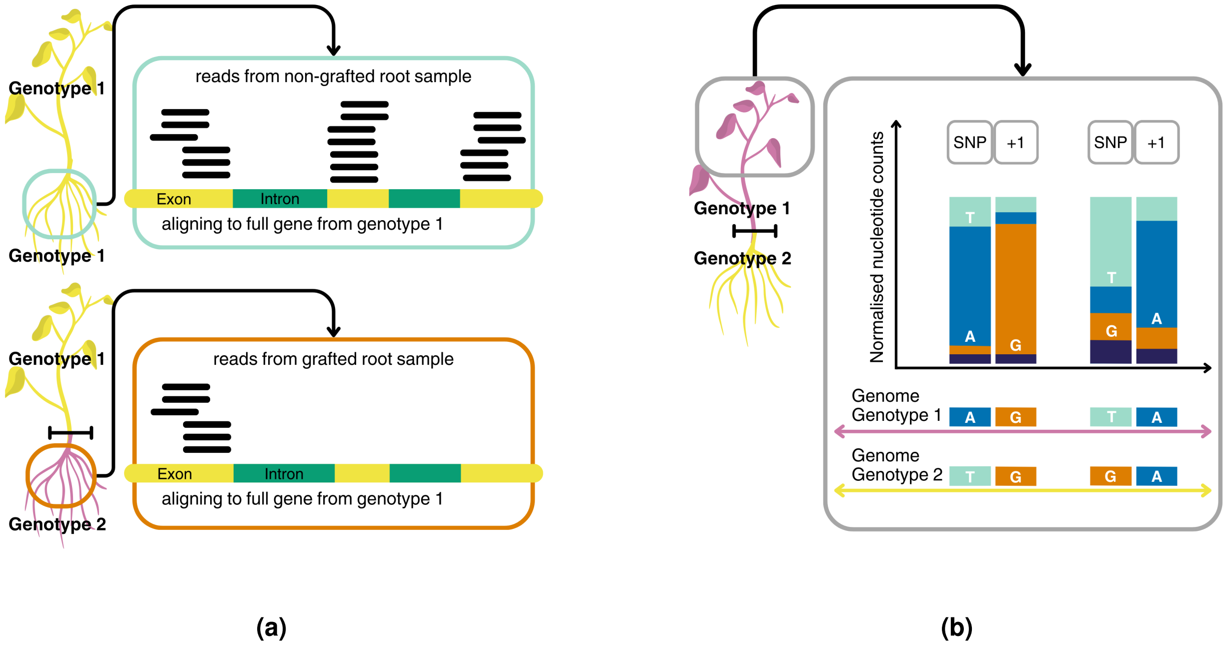


**Supplemental Fig. S9.** Full-length transcript coverage and differences in the distribution of nucleotides between SNPs and other positions enhance the evidence for the presence of a foreign transcript in the sampled tissue. (a) Sequenced transcripts would ideally have RNA-Seq reads covering most of the sequence, i.e. that all exons of the mRNA are approximately equally covered by sequencing reads (top left). Reads covering all exons in the sample from Genotype 2 provide support for the whole transcript having moved from Genotype 1 to Genotype 2 across the graft junction. Transcripts with coverage only for a subsequence (bottom left) do not support full-length presence of the foreign transcript. (b) Neighbouring positions to SNPs can be used as a negative control to evaluate the strength of the signal at SNP positions. Shown here are neighbouring positions of the identified SNPs at the next nucleotide (SNP position +1). If the neighbouring position shows similar levels of alternative nucleotides as the SNP position, the these are likely sequencing errors, rather than evidence for the alternate allele. If the SNP positions have a different frequency of Genotype 2 allele than the neighbouring position has errors, then there is evidence for the alternate allele. Analysing the frequencies of nucleotides at known SNP positions and their neighbours can aid data interpretation.

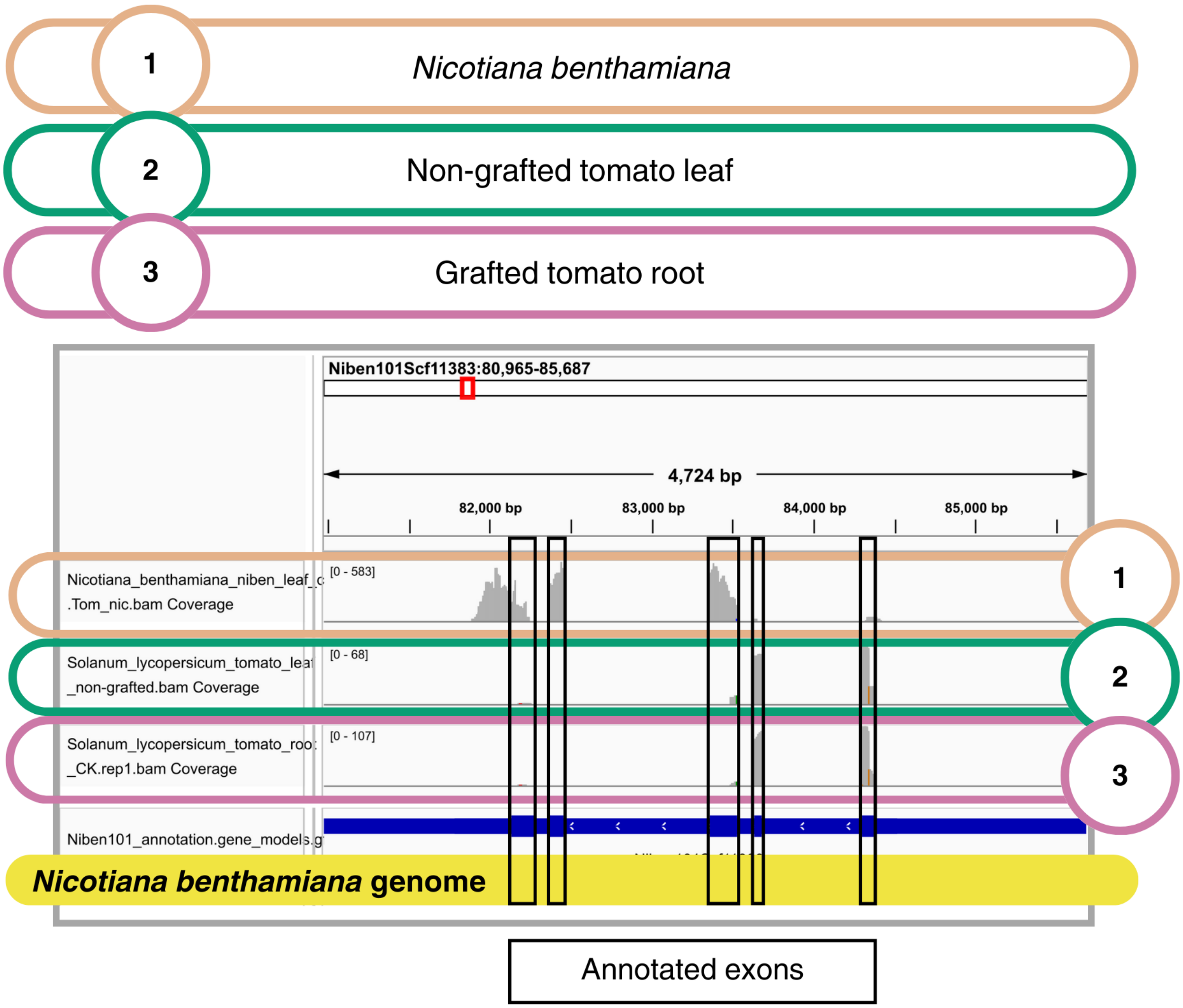


**Supplemental Fig. S10.** An example of poor coverage for a candidate mobile mRNA. In the *Nicotiana benthamiana* annotation of the depicted gene (Niben101Scf11383g00015.1) we find 5 annotated exons of which all are populated with reads at different levels (grey histograms). In the samples from tomato, non-grafted or grafted we can see that (a) not all annotated exons are populated with reads; (b) The exons with coverage are populated in both grafted and non-grafted samples. See Figure 9 and main text for further explanations.This is a screenshot taken in IGV (78).

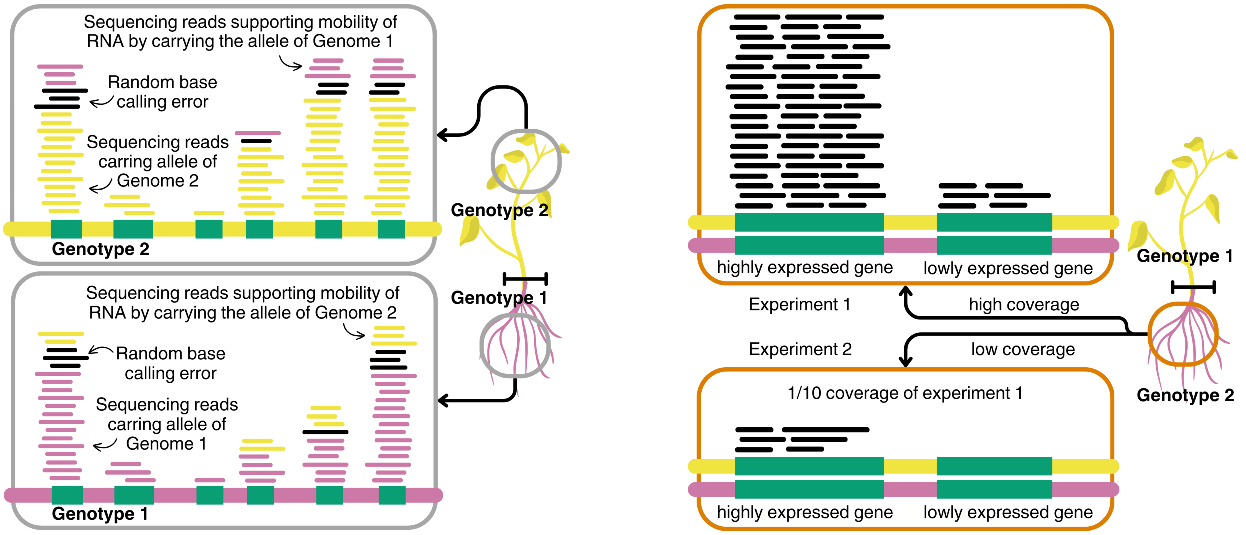


**Supplemental Fig. S11.** Challenges in identifying non-selective mobility versus contamination in high-throughput mobile mRNA detection using RNA-seq data in within-species grafts and cross-species grafts. (a) The presence of Genotype 1 reads in Genotype 2 samples and *vice versa*, across the whole of genome, especially in genes expressed in both tissues is consistent both with non-selective transport and contamination. (b) The two genes presented in this schematic figure have different relative expression levels. In Experiment 2 the sequencing depth is insufficient to detect lowly expressed genes.
